# Alkalinized Filtered Water Induces Changes in the Gut Microbiome in Inflammatory Bowel Disease

**DOI:** 10.1101/2025.09.19.677298

**Authors:** Laura Doblado, Cristina Estebaranz, Elena Carrillo, Alejandro K. Samhan-Arias, Esther Nova, Eva García-Perea, Ascensión Marcos, Ligia Esperanza Díaz, María Monsalve

## Abstract

**Objective:** This study evaluated the effects of filtered, alkalinized water on inflammation and intestinal dysbiosis in individuals with inflammatory bowel disease (IBD).

**Methods:** We conducted a three-month, two-arm, randomized intervention study involving 46 patients with IBD in remission. Participants were divided into two groups: one consumed filtered water from an active filtering device, and the control group consumed water from a mock device. Blood and stool samples were collected before and after the intervention. We assessed antioxidant capacity and circulating cytokine levels from plasma. Gene expression levels of inflammatory mediators were determined using mRNA from peripheral blood mononuclear cells (PBMCs). The microbial composition of fecal samples was characterized by qPCR analysis using primers targeting 16S rRNA genes.

**Results:** Consumption of alkalinized filtered water significantly reduced *IL1B* gene expression in PBMCs. Furthermore, subjects drinking alkalinized filtered water exhibited a consistent, albeit not statistically significant, decrease in circulating IL-1β and significantly lower levels of IL-4 than controls. Microbiome analysis revealed that the levels of the *Bacteroides fragilis* Group were significantly lower in subjects consuming tap water than in those consuming filtered water.

**Conclusion:** These changes suggest that consuming alkalinized water for three months leads to an improved inflammatory status compared with consuming tap water.

## Introduction

Inflammatory bowel disease (IBD), encompassing ulcerative colitis (UC) and Crohn’s disease (CD) is a chronic inflammatory condition of the gastrointestinal tract affecting approximately 6.8 million people worldwide. While genetic susceptibility plays a significant role, IBD primarily arises from the impact of environmental factors on the gut microbiome [1]. For instance, genetic factors explain only ∼13.1% of the variance for CD and 8.2% for UC [2]. The increasing incidence of IBD in newly industrialized countries underscores the influence of a Western lifestyle on risk. Environmental risk factors identified through association studies include: urban living, long-term use of nonsteroidal anti-inflammatory drugs, estrogen-progesterone oral administration (e.g., contraceptives and hormone replacement therapies), antibiotic use, diet regimes high in meat/low in vegetables, and processed food consumption [3]. Current models suggest these risk factors alter the intestinal mucosa immune response to the gut microbiome in genetically susceptible individuals.

Recent advancements in understanding complex mucosa-microbiome interactions and how diet alters microbiome composition suggest that the direct impact of diet on the microbiome is a key factor in IBD development [4]. For example, active IBD is linked to major gut microbiota shifts, including reduced overall diversity, an increase in Proteobacteria (especially Gammaproteobacteria), and a decrease in Firmicutes [5]. Conversely, healthy diet regimes and probiotics have consistently demonstrated the ability to partially reverse this dysbiosis and reduce inflammation [6].

Limited epidemiologic evidence suggests drinking water may be a relevant risk factor for IBD, though the accumulated evidence is still very limited [7]. Evaluating the health effects of tap water consumption is a complex, challenging, and even daunting task. This complexity arises from the varying chemical compositions of tap water, which are influenced by factors such as season, rainfall, soil, and treatment processes [8]. Despite these challenges, several observation studies have explored the links between drinking water, intestinal inflammation, and IBD. Specifically, the presence of iron [9] and other metals, nitrates [10], sulfites, disinfectants, and surfactants [11] [7] have all been associated with intestinal inflammation and IBD incidence. More recently, studies are also examining the contribution of microplastics in water to IBD [12].

Overall, growing health concerns regarding increasing and highly variable pollutant by-products in drinking water (such as metals, microplastics, and persistent organic compounds) have boosted the use of spring/purified bottled water and water filter systems [13]. These alternatives can significantly reduce the concentration of hazardous substances in tap water and can also positively impact the gut microbiome[14].

Water filtering systems also known as point-of-use water filters, can be grouped into four basic categories: 1) particle filters, which remove solid particles but do not alter the water chemistry; 2) reverse osmosis systems, which reduce the total concentration of ions; 3) activated charcoal filters, which are particularly effective in removing chlorine (Cl^-^) and organic compounds; and 4) alternative systems, such as alkalinizing water devices.

Alkalinized water has been recommended for patients with gastric acidosis [17], and in Japan, accumulated evidence has led to the approval of electrolyzed alkaline-reduced water (EARW) apparatuses as medical devices. The impact of drinking water pH on the microbiome is also well-established [15], prompting investigations into the potential intestinal benefits of alkalinized water consumption. However, the beneficial effects of EARW consumption remain controversial [16],[17]. Filter-based ion-exchange systems offer an alternative to electrolysis for alkalinizing water, but scientific studies on their health benefits are still in early stages. Our previous research in rodents showed that filter alkaline water consumption improved vascular reactivity and reduced systemic inflammation [18]. We then sought to determine if this anti-inflammatory activity was linked to intestinal effects. To that end, we used lean and obese Zucker rats, a common preclinical model for obesity and diabetes research due to their insulin resistance, glucose intolerance, and metabolic syndrome [19], We treated these rats with either tap water or alkalinized filtered water for three months. Our findings revealed that the filtered water reduced intestinal inflammation and oxidative stress markers, improved the intestinal mucosa status, and positively affected the microbiome profile in stool samples [20]. Given these promising results, we conducted a human intervention to see if alkalinized filtered water could positively impact intestinal dysbiosis and inflammation in patients with IBD. After three months of alkalinized water use, we observed changes indicative of an improved inflammatory profile compare with subjects who drank tap water.

## Materials and Methods

### Human subjects

This pilot, longitudinal, prospective, randomized, and parallel nutritional intervention study involved two groups. We advertised the study *via* social media, and interested individuals were fully informed about the study. In total, 52 volunteers signed the informed consent and 46 completed the study. The main inclusion criteria were to be an adult patient with IBD with a self-reported remission phase who could provide informed consent. Exclusion criteria included comorbidities and confounding pharmacological treatments. Demographic information was collected from all patients and they all completed a questionnaire on their health, diet, and physical activity. Volunteers were asked to drink filtered water for three months, and received filtering jars containing either active filters (Alkanatur® Drops SLU, Milladoiro-Ames, A Coruña, Spain) or mock filters (Tap). An authorized laboratory (Oliver Rodés) analyzed the water filtering system, confirming its compliance with UNE 149101:2015 standards for human consumption. This system has been shown to remove trihalomethanes and Cl^-^, while adding 15 mg/L of Mg^2+^; it also has been shown to provide water free of microorganisms, and not to release sodium (Na^+^). The plastic jar was approved for food contact and demonstrated to be free of bisphenol A, epoxidized soybean oil, and phthalate esters [21].

A professional nurse collected blood and stool samples at the School of Medicine of the UAM before and after the study. Fasting blood samples were drawn from the cubital vein into BD Vacutainer tubes with EDTA (Becton Dickinson, San Jose, CA; Cat. No. 368589) early in the morning. These samples were used to isolate plasma and PBMCs using density-gradient Ficoll-Paque-PLUS separation (Cytiva, Marlborough, MA; Cat. No.11768538). Stool samples were collected in sterile containers (Acofarma, Madrid, Spain; Cat. No. 311218), then frozen and stored at -80ºC until processing. The study received approval from the Ethics Committees of the IIBM, the UAM, and the Spanish Research Council (*Consejo Superior de Investigaciones Científicas*, CSIC).

### Microbiome

DNA was purified from collected stool samples using the QIAamp® Fast DNA Stool Mini Kit (Quiagen, Cat. No.: 51604). Bacterial quantification was performed by qPCR using primers specific to the bacterial groups: *Bacteroides Fragilis* Group, *Blautia coccoides-Eubacterium rectale* Group, *Clostridium* cluster IV, *Bifidobacterium* spp., *Lactobacillus* spp., *Enterobacteriaceae, Enterococcus* spp., *Faecaibacterium praustnizii* and *Akkermansia muciniphila* directed against 16S rRNA coding genes [22][23][24][25][26][27].

### Gene expression analysis

PBMCs were lysed in the presence of 1 mL of TRIzol™ reagent (ThermoFisher Scientific, Waltham, MA; Cat. No.: 15596018) and total RNA was isolated following the manufacturer’s instructions. cDNA was synthesized from total RNA preparations by reverse transcription of 1 µg of RNA using MMV reverse transcriptase (Promega Biotech, Madison, WI; Cat. No.: M1701), in a final volume of 20 µL. The mixture was incubated at 37°C for 45 min and then cooled for 2 min at 4°C. The resulting cDNA then served as the template for subsequent qPCR. The primers used are listed below. Each 10 µL PCR reaction included 1 µL cDNA, 5 µL qPCRBIO SyGreen Mastermix (Cultek SL, Dutchcher Group, Madrid, Spain; Cat. No.: PB20.14-01) and primers (0.3 µM). Samples were analyzed in triplicate on a Mastercycler® RealPlex2 (Eppendorf,Hamburg, Germany) using *36B4* as the loading control.

*36B4*

Fw 5’-GCGACCTGGAGTCCAACTA-3’

Rv 5’-ATCTGCTGCATCTGCTTGG-3’

*IL1B*

Fw 5’-GAAGCTGATGGCCCTAAACAG-3’

Rv 5’-AGCATCTTCCTCAGCTTGTCC-3’

*IL4*

Fw 5’-*GCTGCCTCCAAGAACACAAC*-3’

Rv 5’-*TCACAGGACAGGAATTCAAG*-3’

*IL6*

Fw 5’-GGCACTGGCAGAAAACAACC-3’

Rw 5’-GCAAGTCTCCTCATTGAATCC-3’

*IL10*

Fw 5’-ACCTGCCTAACATGCTTCGAG-3’

Rw 5’-CTGGGTCTTGGTTCTCAGCTT-3’

*TGFB1*

Fw 5’-GAGCCTGAGGCCGACTACTA-3’

Rv 5’-CGGAGCTCTGATGTGTTGAA-3’

*TNF*

Fw 5’-GTGCTTGTTCCTCAGCCTCTT-3’

Rv 5’-ATGGGCTACAGGCTTGTCACT-3’

*IFNG*

Fw 5’-GAATAACTATTTTAACTCAAGTGGCA-3’

Rv 5’-CAGGATTTTCATGTCACCATCCTT-3’

### Cytokines

Circulating levels of IL-1β, IL-4, IL-6, TNF-α, and IL-10 in plasma were analyzed by cellular cytometry using Multiplex Cytokine Assays (Pro-cartaPlex Immunoassays, ThermoFisher Scientific) at the CNB (CSIC) Flow Cytometry Unit.

### Antioxidant capacity

Antioxidant capacity was determined in plasma samples using the e-BQC electrochemical analytical system (BioQuoChem, Oviedo, Spain), which measures total (QT), fast (Q1) and slow-acting (Q2) antioxidant capacity.

### Statistical analysis

Microsoft Excel (Microsoft Corp., Redmond, WA) was used for data processing. GraphPad Prism 9 (GraphPad Software Inc., San Diego, CA) was used for statistical analysis and graph preparation. The normality of the data was assessed using the Kolmogorov-Smirnov test. Statistical significance of differences between groups was assessed using two-tailed *t*-test, and two-way ANOVA for continuous variables. Levene’s test was used for equality of variances. Fisher’s exact test was used for discontinuous variables. The values were considered statistically significant when *p* < 0.05. No values were discarded; those values not included were due to failure in sample acquisition or technical problems.

## Results

To assess the effects of alkalinized filtered water in patients with IBD, we conducted a three-month intervention study. Fifty-two volunteers signed informed consent forms. We randomly assigned 26 participants to receive a mock filter unit and 26 to receive an active filter unit (Table 1). Participants were asked to complete a form with epidemiological data and use their assigned unit for all drinking water over the study period. Blood and stool samples were collected prior to the initiation of the study and following the three-month intervention. Participants received no compensation. Ultimately, 46 subjects completed the study: 25 used the active filtering unit and 21 used the mock filtering unit. Therefore, only one person from the active filter group dropped compared with five from the mock filter group.

**Table 1.** Base line cohort characteristics. Crohn’s disease (CD) and ulcerative colitis (UC).

|  | Tap<br>(n=21) | SD | Filter<br>(n=26) | SD | *p* |
| --- | --- | --- | --- | --- | --- |
| Drop-out | 5 |  | 1 |  | 0.19 |
| Males | 5 |  | 8 |  | 0.74 |
| Females | 16 |  | 17 |  |  |
| Age (year) | 41.24 | 8.85 | 38.76 | 11.16 | 0.41 |
| Weight (kg) | 65.24 | 14.68 | 65.01 | 12.59 | 0.96 |
| Body Mass Index (BMI) | 24.02 | 4.38 | 23.14 | 3.58 | 0.47 |
| Crohn's Disease (CD) | 14 |  | 14 |  | 0.76 |
| Ulcerative Colitis (UC) | 5 |  | 12 |  |  |

### Circulating cytokines

Following the intervention, the levels of IL-4 were significanlty lower in the filtered water group than in the tap water group, along with a trend for reduced IL-1β levels (*p* = 0.05). However, no significant differences were found when comparing the two intervention groups (Fig. 1). Overall, neither group experienced an increase in any of the tested cytokines over time. While the observed reductions were not statistically significant, the filtered water group generally showed slightly greater reductions for all cytokines with the exception of IL-10 (Fig. 2). This suggests that drinking alkalinized filtered water may have a mild inmunomodulatory effect.

**Figure 1.**
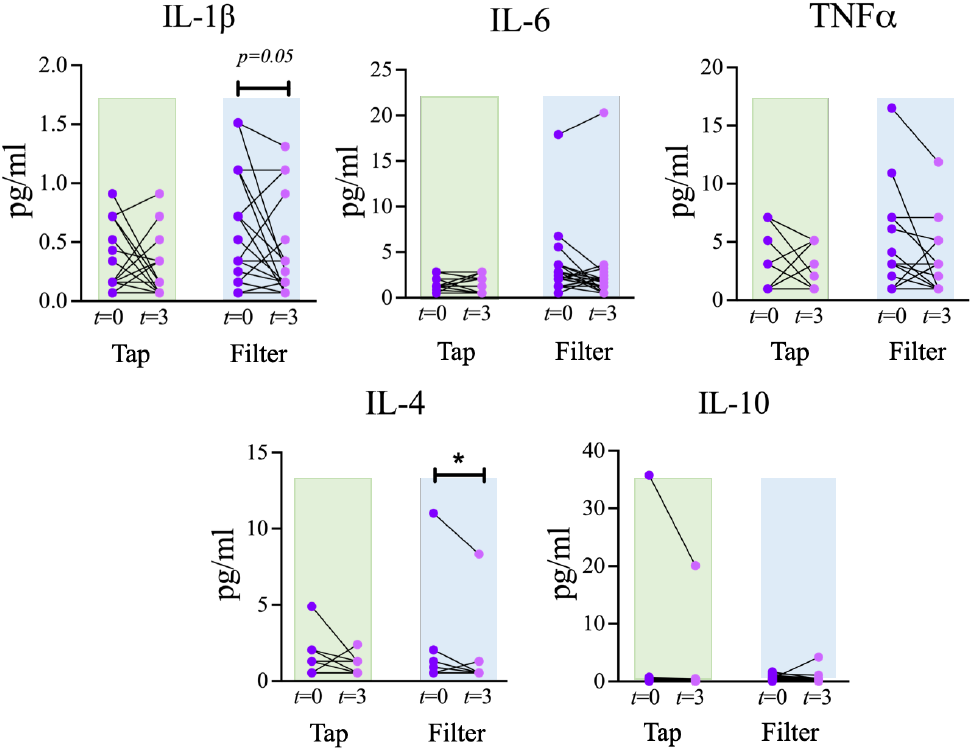
Analysis of the indicated cytokines in plasma blood samples from volunteers before and after the intervention dinking tap or filtered water. The graphs show individual values in pg/mL.\**p*<0.05 (*t* test).

**Figure 2.**
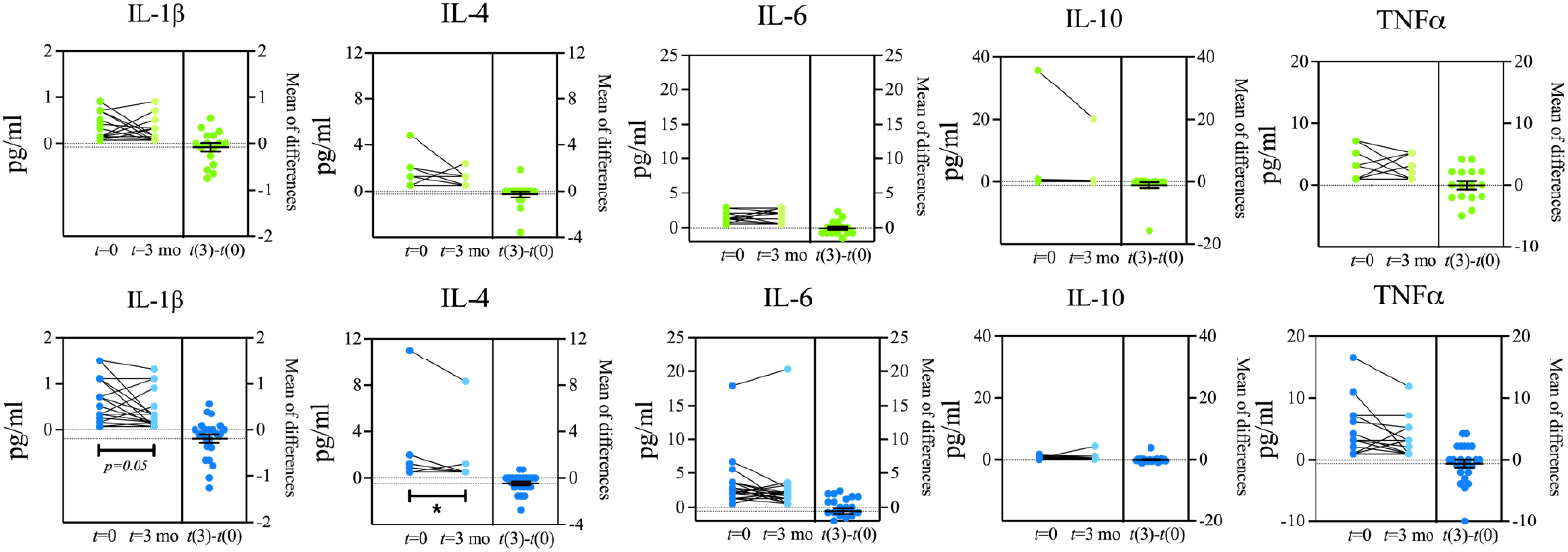
Analysis of the indicated cytokines in plasma samples from volunteers before and after the intervention dinking tap or filtered water. The left section of the graphs shows the change in individual values at *t*=0 and *t*=3 months in pg/mL. The right section of the graph shows the variation of the values for each volunteer with the intervention and includes mean +/-SEM.

### Analysis of cytokine gene expression

The analysis of circulating cytokines was complemented with a gene expression analysis in PBMCs: *IL1B* (encoding IL-1β), *IL*4, *IL6, IL-10, TGFB1* (encoding TGFβ), *TNF* (encoding TNF α) and *IFNG* (encoding IFNγ). Following the intervention, the mRNA levels of *IL1B* were significantly higher in the tap water group, while the filtered water group exhibited no such response. This led to significant differences in *IL1B* expression between the groups after the intervention. No other significant changes were identified in the remaining genes tested, either between groups or as a result of the intervention (Fig. 3). Overall, in the tap water group gene expression levels did not change with the intervention, while in the filtered water group, gene expression levels on average decreased or did not change. Nevertheless, only the differences in *IL1B* expression reached statistical significance (Fig. 3, 4).

**Figure 3.**
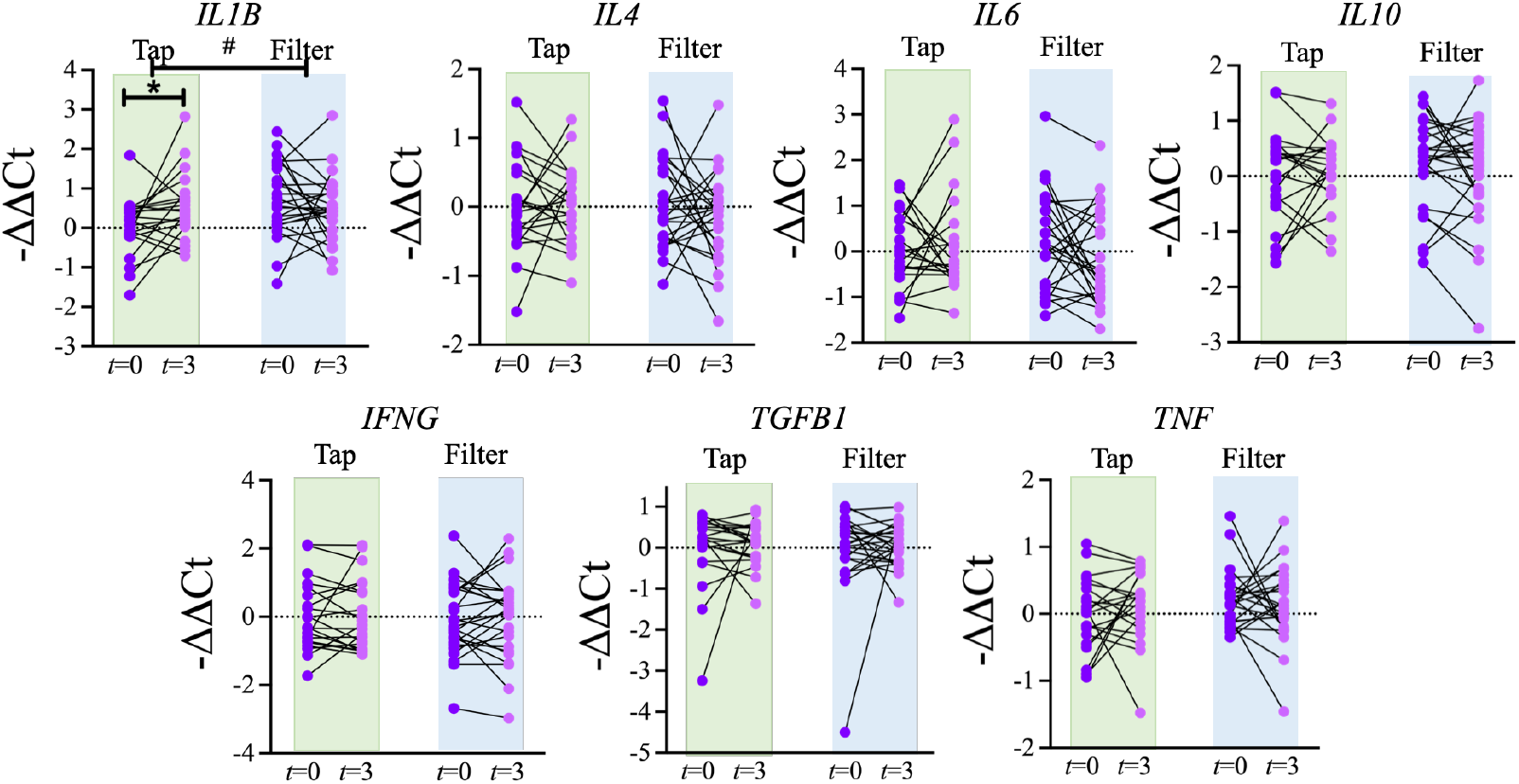
qRT-PCR gene expression analysis of the indicated cytokine-coding genes in PBMCs. The graphs show -1^′^1^′^Ct individual values, *36B4* was used as a loading control and data was normalized to the mean of the tap water group prior to the intervention. \**p*<0.05 (*t* test). # *p*<0.05 (two-way ANOVA).

**Figure 4.**
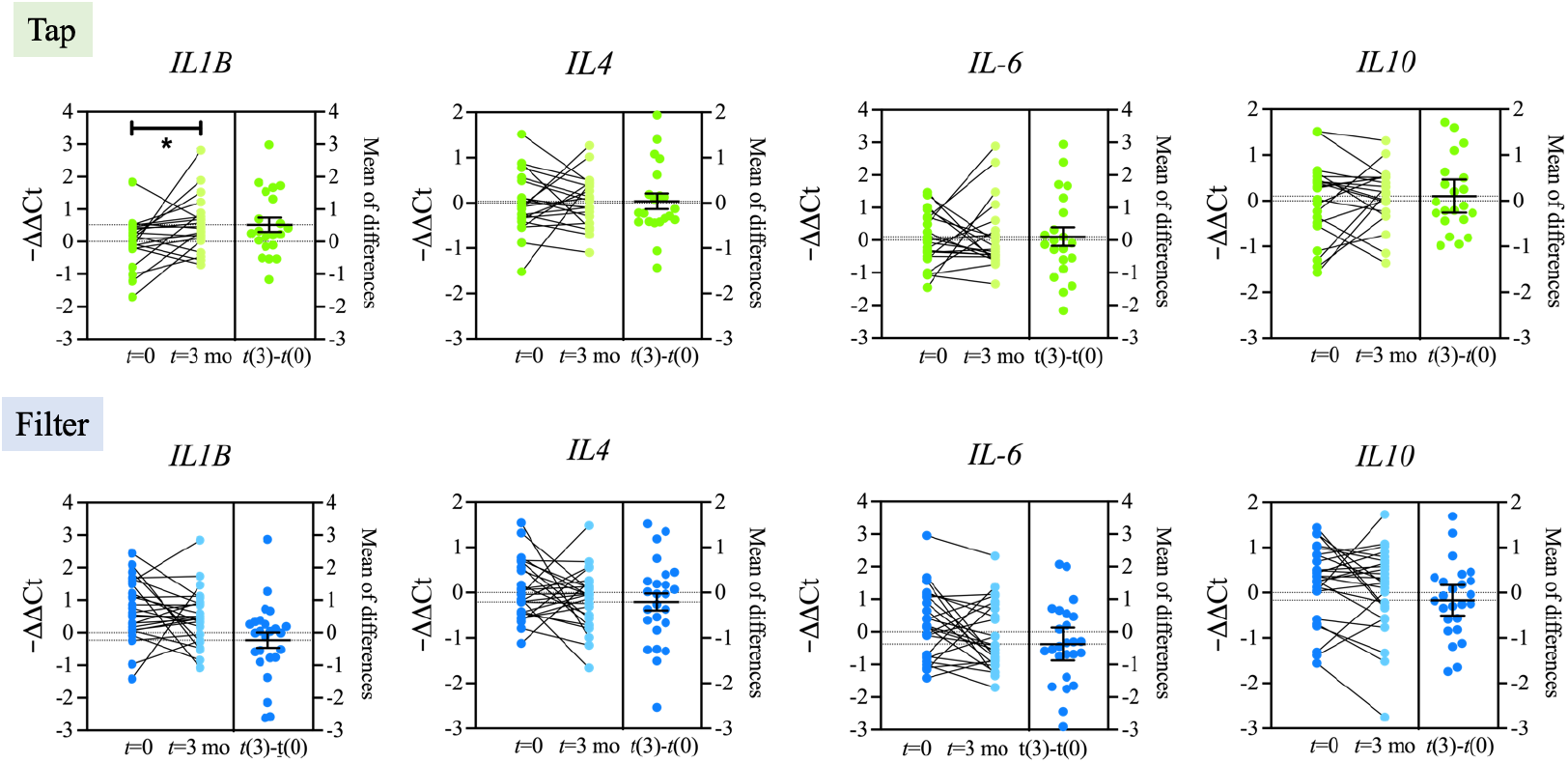
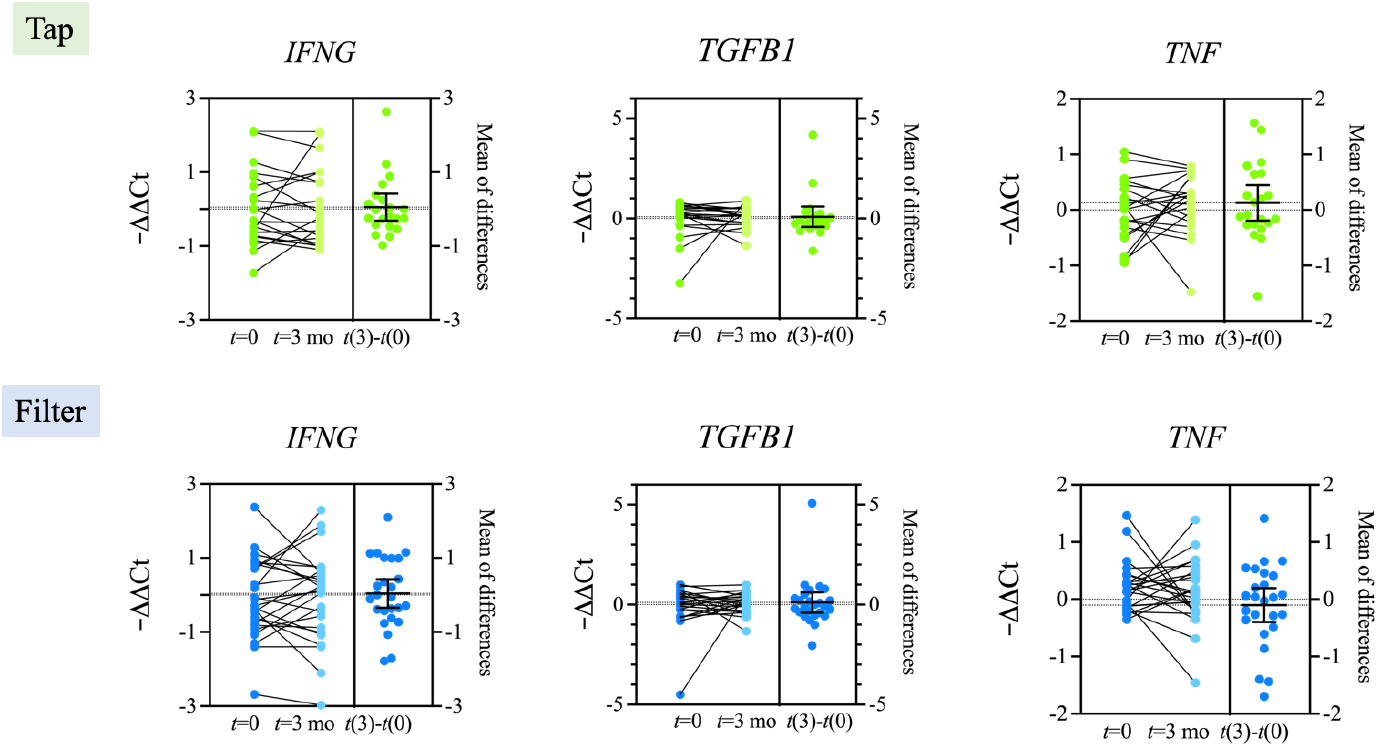
qRT-PCR gene expression analysis of the indicated cytokine-coding genes in PBMCs. The left section of the graphs shows the change in the individual -ΔΔCt values at *t*=0 and *t*=3 months in pg/ml. *36B4* was used as a loading control and data normalized to the mean of the tap water group prior to the intervention. \**p*<0.05 (*t* test). # *p*<0.05 (two-way ANOVA). The right section of the graph shows the variation of the values for each patient with the intervention and includes mean +/-SEM.

### Antioxidant capacity

We next evaluated the impact of the intervention on plasma antioxidant capacity. Accordingly, fast (Q1), slow (Q2), and total (QT) antioxidant capacities were measured using the e-BQC electrochemical reader. No significant differences were observed either within or between the groups (Fig. 5, 6), suggesting that the water source did not affect the plasma antioxidant capacity of the volunteers. Overall, both intervention groups showed a non-significant trend toward reduced antioxidant capacity. Notably, the inter-individual variability in antioxidant capacity was non-significantly more reduced in the tap water group than in the filtered water group. A reduced antioxidant capacity might relate to a decreased inflammatory state, an induced compensatory antioxidant response, or a dimished systemic capacity to manage pro-oxidants. In the present context, tap water appeared to be more pro-inflamatory, while filtered water seemed more anti-inflamatory. The subtle changes observed could point to a better redox balance in individuals who drank filtered water. However, the lack of statistical significance limits the findings robustness, likely due to the small sample size.

**Figure 5.**
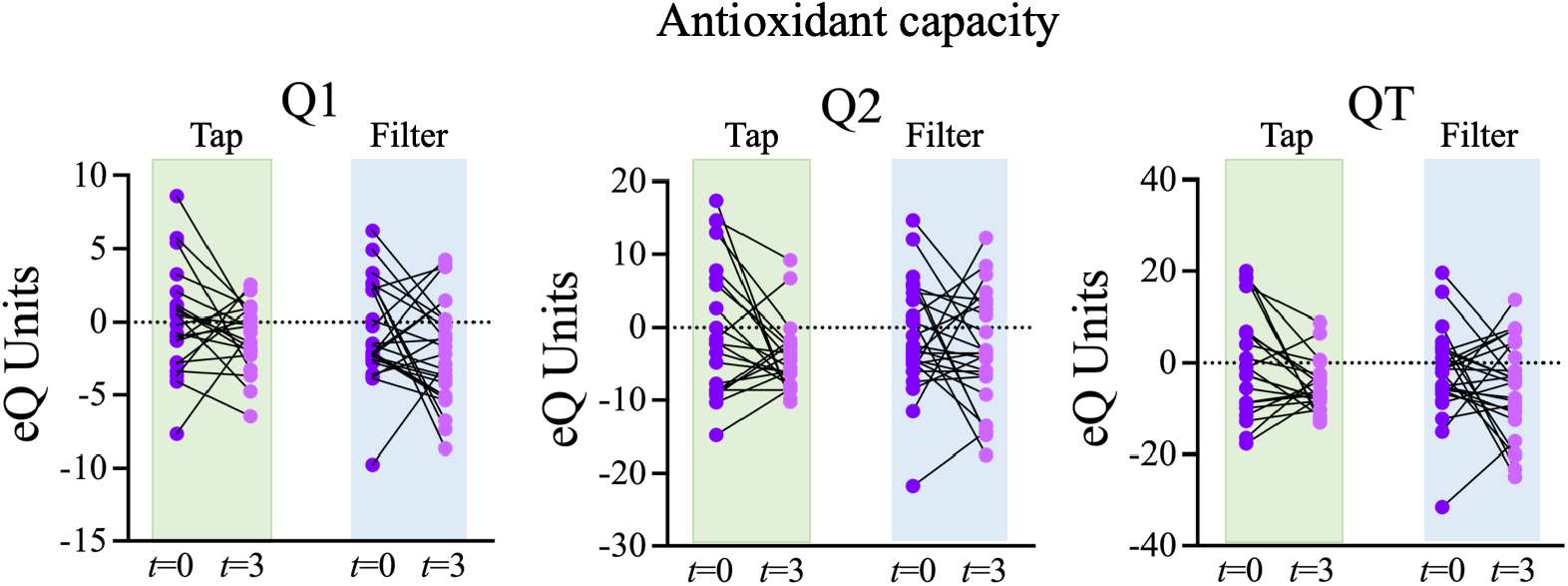
Analysis of the fast (Q1), slow (Q2) and total (QT) antioxidant capacities in plasma samples from volunteers before and after the intervention drinking tap or filtered water. The graphs show the individual values difference with the average of the tap group at *t*=0 in electrochemical units.

**Figure 6.**
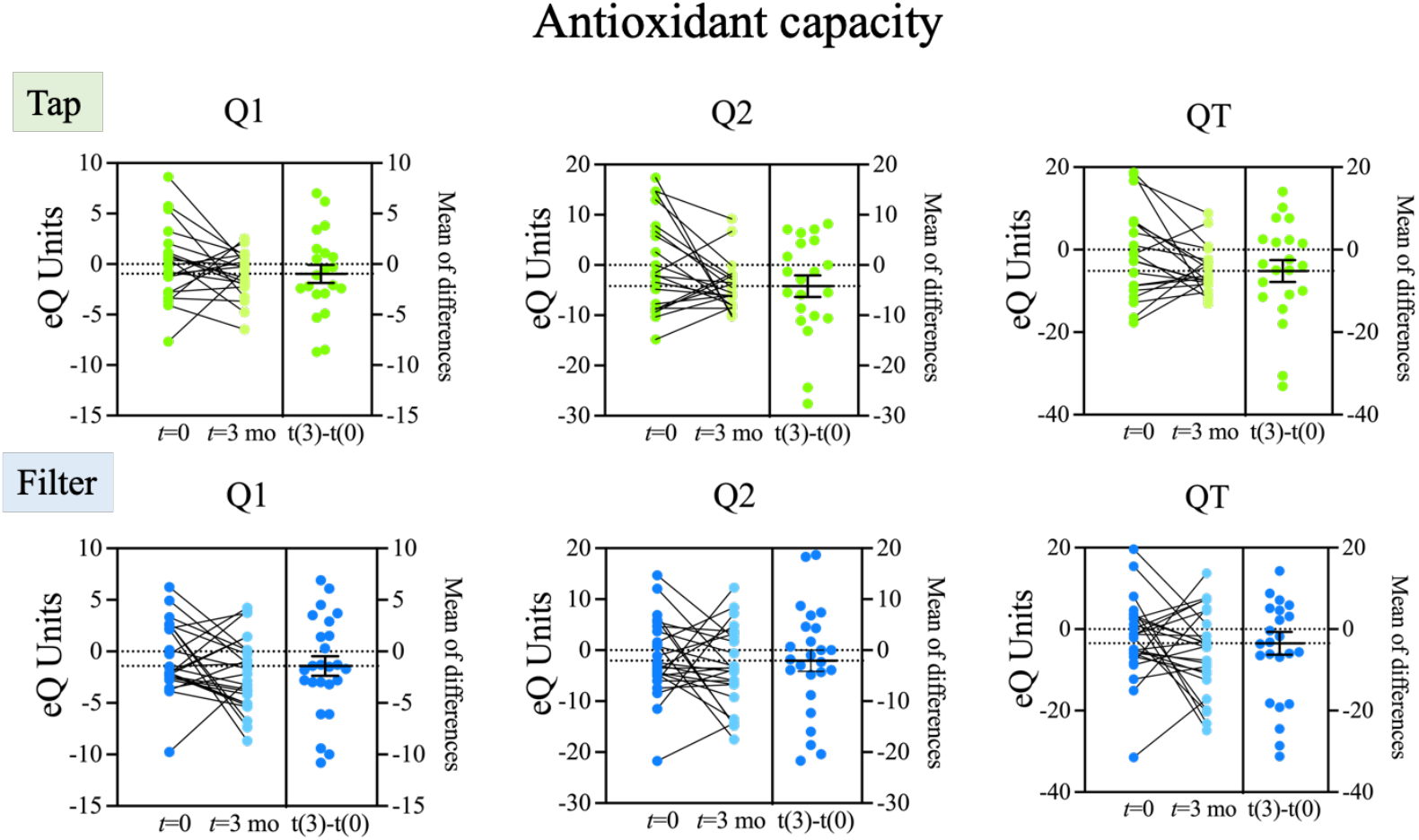
Analysis of the fast (Q1), slow (Q2) and total (QT) antioxidant capacity in plasma samples from volunteers before and after the intervention drinking tap or filtered water. The left section of the graphs shows individual value differences with the average of the group at *t*=0 in electrochemical units. The right section of the graph shows the variation of the values for each participant with the intervention and includes mean +/-SEM.

### Intestinal microbiome

Finally, we evaluated the impact of the intervention on the intestinal microbial composition. To do this, total DNA was isolated from stool samples collected before and after the intervention, and was analyzed by qPCR. We assessed changes in nine bacterial groups previously linked to metabolic status or diet [28][29][30]: *Bacteroides fragilis* Group, *Blautia coccoides-Eubacterium rectale* Group, *Clostridium* cluster IV, *Bifidobacterium* spp., *Lactobacillus* spp., *Enterobacteriaceae, Enterococcus* spp., *Faecaibacterium praustnizii* and *Akkermansia muciniphila*. In samples prior to intervention (t=0) we found a significant difference was found among the groups. The levels of *Bifidobacterium* spp. were significantly lower in the filtered water group than in the tap water group. This unpredictable bias, despite randomized device distribution, is notable because reduced *Bifidobacterium* spp. have been found in patients with bowel disease and are associated with metabolic alterations such as obesity [31]. In response to the intervention, we found a significant reduction in the *Bacteroides fragilis* Group in the tap water group, while this effect was not observed in the filtered water group. However, no statistically significant differences were found between both groups (Fig. 7). The *Bacteroides fragilis* Group has been extensively studied, includes common anaerobe members of the human microbiota and has been shown to provide several immunomodulatory benefits to healthy hosts [32]. Its role in IBD remains controversial. Some enterotoxic strains are elevated in IBD and associated with the active phase of the disease [33]. Conversely, the *Bacteroides fragilis* Group has also been shown to alleviate the disease when used as a probiotic because of its anti-inflammatory effects associated with IL-10 induction [34]. Similarly, in other diseases such as obesity, accumulated data suggest that these bacteria can behave differently depending on the context, playing both pro- and anti-inflammatory roles [35]. A relevant study that shed some light on this context-dependent behavior of *Bacteroides fragilis* Group showed that these bacteria regulate enterotoxin gene expession by altering the orientation of the promoter of the enterotoxin gene. This modification is increased in IBD and linked to its pro-inflamatory role, suggesting that such pathological promoter modification could result in a pro-inflammatory phenotype [36].

**Figure 7.**
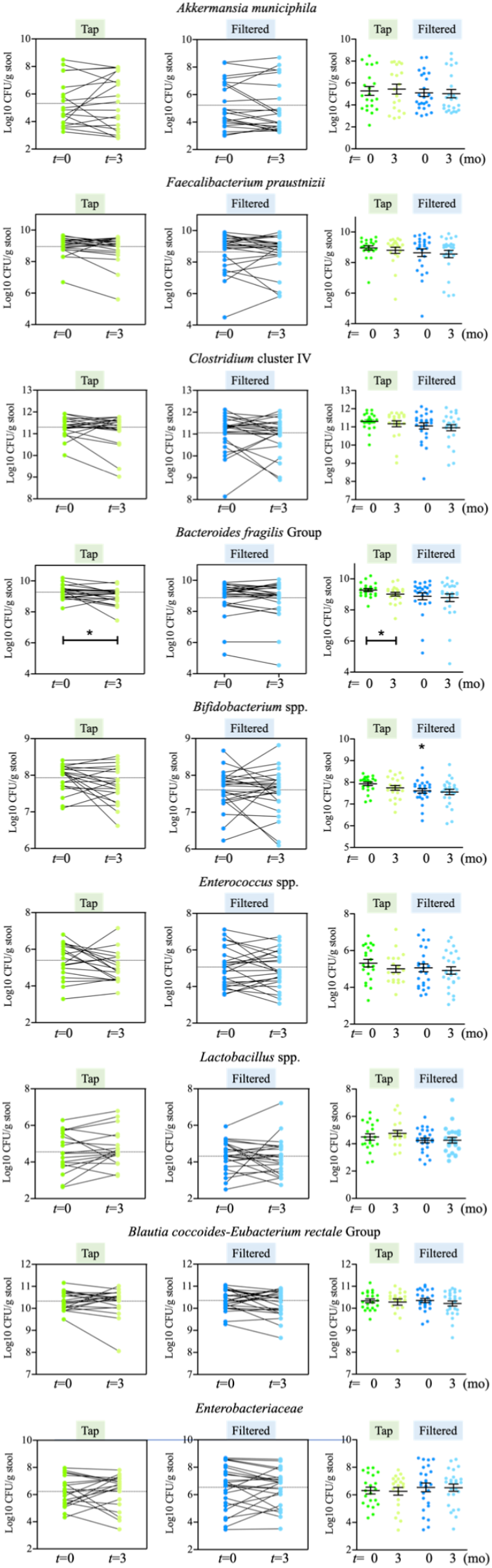
qPCR analysis of fecal microbial content. The graphs on the left panels show how individual values changed according to the intervention. The right panels show the distribution of individual values prior to and after intervention, and include mean +/-SEM. Statistical comparison between groups was carried out using *t* test type 1 (variations over time) and type 2 (differences between groups). \**p*<0.05.

## Discussion

The rising prevalence of IBD has intensified the search for risk factors and complementary interventions to traditional medications. Current evidence suggests that in genetically susceptible individuals, dietary factors can contribute to disease development. Moreover, diet-associated dysbiosis is a significant component of IBD, and some probiotics have demonstrated positive immunosuppressive effects in this context. Water, though often overlooked, is a vital dietary component. In fact, water quality has been identified as a risk factor for IBD, implying that consuming the right kind of water might also be relevant in disease development.

Our prior preclinical studies in mice and rats revealed that consuming filtered water from an alkalinizing device for three months reduced systemic and intestinal inflammation and improved intestinal dysbiosis compared with tap water. These findings, based on targeted microbiome analysis of feces [18][20] led us to investigate similar effects in humans with IBD. We evaluated a cohort of 46 volunteers, analyzing blood samples for inflammatory markers. The results indicated significant or near-significant trends suggesting improved inflammatory status in individuals using the alkalinizing filter device. Notably, these changes were consistent with our preclinical observations. Specifically, after three months, we observed an almost significant (*p* = 0.05) reduction in circulating IL-1ß levels in subjects who consumed filtered water, a change not seen in the tap water group. Further supporting these findings, gene expression analysis in PBMCs showed that *IL1B* (coding for IL-1ß) gene expression increased in tap water consumers after three months, while filtered water consumers exhibited, on average, lower levels. The interaction between tap and filtered water consumption was statistically significant. These results align with our previous findings demonstrating that filtered water consumption reduced circulating IL-1ß levels in lean Zucker rats and in SHR rats after a three-month intervention [18][20]. Collectively, these results suggest that consuming filtered water is associated with lower IL-1ß levels compared with tap water.

These findings are particularly relevant given the extensive research supporting IL-1β as a crucial mediator of inflammation and tissue damage in IBD. IL-1β significantly contributes to pro-inflammatory responses by recruiting and activating immune cells in the gut mucosa. Moreover, it is involved in disrupting the intestinal barrier and modulating the differentiation and function of T helper cells, which are known to play a role in IBD development. Notably, intestinal dysbiosis has been shown to stimulate IL-1β release, thereby promoting inflammation [37].

Since inflammation is commonly associated with oxidative stress, and our previous rat data suggested that filtered water reduced levels of with 4-HNE-modified proteins (an indicator of oxidative stress), we evaluated changes in plasma oxidative capacity. However, no significant differences were found between the groups, likely due to the small sample size.

We conducted a targeted analysis of bacterial content in stool samples. Our prior research in Zucker rats demonstrated that filtered water increased the abundance of *Akkermansia muciniphila* and decreased *Enterobacteriaceae* levels, leading to an improved microbiome profile [20]. However, we did not observe the same changes in the IBD cohort of the current study. The only significant difference linked to the intervention was a reduction in the *Bacteroides fragilis* group in the group that consumed tap water; this effect was not seen in the group who consumed filtered water. Interestingly, our previous study with lean Zucker rats also showed that tap water consumption reduced the abundance of the *Bacteroides fragilis* group while filtered water had the opposite effect [20]. This consistency suggests the outcome is likely functionally significant. Interpreting these findings requires considering the inflammatory status. The *Bacteroides fragilis*

Group typically has an anti-inflammatory role. However, during the active phase of IBD, these bacteria can activate an enterotoxin known to negatively impact disease progression. Given the relatively higher levels of IL-1ß observed in the tap water group (in both rats and humans), it is plausible that the reduction of the *Bacteroides fragilis* group in tap water consumers is linked to their increased inflammation.

Overall, these data support the hypothesis that consuming alkalinized filtered water may improve the inflammatory state in patients with IBD and potentially reduce dysbiosis. However, due to the limited number of subjects and the specific biomarkers evaluated, further research is needed to confirm the significance of these findings.

## Author Contributions

conceptualization, M.M. & A.S; methodology, L.E.D., E.N., A.M., M.M.; formal analysis, M.M. & L.E.D.; investigation, L.D., L.E.D.; resources, M.M.; data curation, M.M. & L.E.D.; writing—original draft preparation, M.M. & L.D.; writing—review and editing, M.M.; supervision, A.M., L.E.D., M.M.; project administration, A.M., M.M.; funding acquisition, M.M. All authors have read and agreed to the published version of the manuscript.

## Data Availability Statement

Data supporting reported results is included as Supplementary Material.

## Acknowledgments

Technical support was provided by Raquel García. Editorial support was provided by Dr. Kenneth McCreath. Our special thanks to ACCU ESPAÑA and ACCU Madrid for supporting and guiding the preparation of the study, the social media advertisements and its distribution as well for providing the necessary framework for the registration of volunteers and collection of demographic data.

## References

1. Ananthakrishnan, A.N. Epidemiology and Risk Factors for IBD. Nat Rev Gastroenterol Hepatol 2015, 12, 217–, doi:10.1038/nrgastro.2015.34.

2. Liu, J.Z.; van Sommeren, S.; Huang, H.; Ng, S.C.; Alberts, R.; Takahashi, A.; Ripke, S.; Lee, J.C.; Jostins, L.; Shah, T.; et al. Association Analyses Identify 38 Susceptibility Loci for Inflammatory Bowel Disease and Highlight Shared Genetic Risk across Populations. Nat Genet 2015, 47, 986–, doi:10.1038/ng.3359.

3. Caron, B.; Honap, S.; Peyrin-Biroulet, L. Epidemiology of Inflammatory Bowel Disease across the Ages in the Era of Advanced Therapies. Journal of Crohn’s and Colitis 2024, 18, ii3–ii15, doi:10.1093/ecco-jcc/jjae082.

4. Zsálig, D.; Berta, A.; Tóth, V.; Szabó, Z.; Simon, K.; Figler, M.; Pusztafalvi, H.; Polyák, É. A Review of the Relationship between Gut Microbiome and Obesity. Applied Sciences 2023, 13, 610, doi:10.3390/app13010610.

5. Lee, M.; Chang, E.B. Inflammatory Bowel Diseases (IBD) and the Microbiome—Searching the Crime Scene for Clues. Gastroenterology 2021, 160, 537–, doi:10.1053/j.gastro.2020.09.056.

6. Roy, S.; Dhaneshwar, S. Role of Prebiotics, Probiotics, and Synbiotics in Management of Inflammatory Bowel Disease: Current Perspectives. World Journal of Gastroenterology 2023, 29, 2100–, doi:10.3748/wjg.v29.i14.2078.

7. Zhou, S.; Chai, P.; Dong, X.; Liang, Z.; Yang, Z.; Li, J.; Teng, G.; Sun, S.; Xu, M.; Zheng, Z.-J.; et al. Drinking Water Quality and Inflammatory Bowel Disease: A Prospective Cohort Study. Environ Sci Pollut Res 2023, 30, 71183–, doi:10.1007/s11356-023-27460-w.

8. Pan, F.; Zhu, S.; Shang, L.; Wang, P.; Liu, L.; Liu, J. Assessment of Drinking Water Quality and Health Risk Using Water Quality Index and Multiple Computational Models: A Case Study of Yangtze River in Suburban Areas of Wuhan, Central China, from 2016 to 2021. Environ Sci Pollut Res 2024, 31, 22758–, doi:10.1007/s11356-024-32187-3.

9. Aamodt, G.; Bukholm, G.; Jahnsen, J.; Moum, B.; Vatn, M.H.; the IBSEN Study Group The Association Between Water Supply and Inflammatory Bowel Disease Based on a 1990–1993 Cohort Study in Southeastern Norway. American Journal of Epidemiology 2008, 168, 1072–, doi:10.1093/aje/kwn218.

10. Holik, D.; Bezdan, A.; Marković, M.; Orkić, Ž.; Milostić-Srb, A.; Mikšić, Š.; Včev, A. The Association between Drinking Water Quality and Inflammatory Bowel Disease-A Study in Eastern Croatia. Int J Environ Res Public Health 2020, 17, 8495, doi:10.3390/ijerph17228495.

11. Xu, Y.; Li, Y.; Scott, K.; Lindh, C.H.; Jakobsson, K.; Fletcher, T.; Ohlsson, B.; Andersson, E.M. Inflammatory Bowel Disease and Biomarkers of Gut Inflammation and Permeability in a Community with High Exposure to Perfluoroalkyl Substances through Drinking Water. Environmental Research 2020, 181, 108923, doi:10.1016/j.envres.2019.108923.

12. Ji, J.; Wu, X.; Li, X.; Zhu, Y. Effects of Microplastics in Aquatic Environments on Inflammatory Bowel Disease. Environmental Research 2023, 229, 115974, doi:10.1016/j.envres.2023.115974.

13. Li, L.-Y.; Duan, Y.-J.; Hou, J. Research Progress on Source, Risk Assessment, and Management of Emerging Pollutants in Drinking Water. Ying Yong Sheng Tai Xue Bao 2023, 34, 3456–, doi:10.13287/j.1001-9332.202312.029.

14. Vanhaecke, T.; Bretin, O.; Poirel, M.; Tap, J. Drinking Water Source and Intake Are Associated with Distinct Gut Microbiota Signatures in US and UK Populations. The Journal of Nutrition 2022, 152, 182–, doi:10.1093/jn/nxab312.

15. Hansen, T.H.; Thomassen, M.T.; Madsen, M.L.; Kern, T.; Bak, E.G.; Kashani, A.; Allin, K.H.; Hansen, T.; Pedersen, O. The Effect of Drinking Water pH on the Human Gut Microbiota and Glucose Regulation: Results of a Randomized Controlled Cross-over Intervention. Sci Rep 2018, 8, 16626, doi:10.1038/s41598-018-34761-5.

16. Higashimura, Y.; Baba, Y.; Inoue, R.; Takagi, T.; Uchiyama, K.; Mizushima, K.; Hirai, Y.; Ushiroda, C.; Tanaka, Y.; Naito, Y. Effects of Molecular Hydrogen-Dissolved Alkaline Electrolyzed Water on Intestinal Environment in Mice. Med Gas Res 2018, 8, 11–, doi:10.4103/2045-9912.229597.

17. Bajgai, J.; Kim, C.-S.; Rahman, M.H.; Jeong, E.-S.; Jang, H.-Y.; Kim, K.-E.; Choi, J.; Cho, I.-Y.; Lee, K.-J.; Lee, M. Effects of Alkaline-Reduced Water on Gastrointestinal Diseases. Processes 2022, 10, 87, doi:10.3390/pr10010087.

18. García-Gómez, R.; Prieto, I.; Amor, S.; Patel, G.; Fuente, M.; Granado, M.; Monsalve, M. Evaluation of the Potential Benefits of Alkaline Drinking Water on Tumor Development Reveals Vascular Protective Effects. Arch Med Sci Civil Dis 2021, 6, 102–, doi:10.5114/amscd.2021.109241.

19. Melnyk, S.; Hakkak, R. Metabolic Status of Lean and Obese Zucker Rats Based on Untargeted and Targeted Metabolomics Analysis of Serum. Biomedicines 2022, 10, 153, doi:10.3390/biomedicines10010153.

20. Doblado, L.; Díaz, L.E.; Nova, E.; Marcos, A.; Monsalve, M. Intestinal Effects of Filtered Alkalinized Water in Lean and Obese Zucker Rats. Microorganisms 2024, 12, 316, doi:10.3390/microorganisms12020316.

21. Certificados Alkanatur. Productos avalados por profesionales Available online: https://alkanatur.com/es/certificados/ (accessed on 22 December 2023).

22. Bartosch, S.; Fite, A.; Macfarlane, G.T.; McMurdo, M.E.T. Characterization of Bacterial Communities in Feces from Healthy Elderly Volunteers and Hospitalized Elderly Patients by Using Real-Time PCR and Effects of Antibiotic Treatment on the Fecal Microbiota. Appl Environ Microbiol 2004, 70, 3581–, doi:10.1128/AEM.70.6.3575-3581.2004.

23. Matsuki, T.; Watanabe, K.; Fujimoto, J.; Kado, Y.; Takada, T.; Matsumoto, K.; Tanaka, R. Quantitative PCR with 16S rRNA-Gene-Targeted Species-Specific Primers for Analysis of Human Intestinal Bifidobacteria. Appl Environ Microbiol 2004, 70, 173–, doi:10.1128/AEM.70.1.167-173.2004.

24. Rinttilä, T.; Kassinen, A.; Malinen, E.; Krogius, L.; Palva, A. Development of an Extensive Set of 16S rDNA-targeted Primers for Quantification of Pathogenic and Indigenous Bacteria in Faecal Samples by Real-time PCR. Journal of Applied Microbiology 2004, 97, 1177–, doi:10.1111/j.1365-2672.2004.02409.x.

25. Heilig, H.G.H.J.; Zoetendal, E.G.; Vaughan, E.E.; Marteau, P.; Akkermans, A.D.L.; de Vos, W.M. Molecular Diversity of Lactobacillus Spp. and Other Lactic Acid Bacteria in the Human Intestine as Determined by Specific Amplification of 16S Ribosomal DNA. Applied and Environmental Microbiology 2002, 68, 123–, doi:10.1128/AEM.68.1.114-123.2002.

26. Walter, J.; Hertel, C.; Tannock, G.W.; Lis, C.M.; Munro, K.; Hammes, W.P. Detection of Lactobacillus, Pediococcus, Leuconostoc, and Weissella Species in Human Feces by Using Group-Specific PCR Primers and Denaturing Gradient Gel Electrophoresis. Appl Environ Microbiol 2001, 67, 2585–, doi:10.1128/AEM.67.6.2578-2585.2001.

27. Derrien, M.; Vaughan, E.E.; Plugge, C.M.; de Vos, W.M. Akkermansia Muciniphila Gen. Nov., Sp. Nov., a Human Intestinal Mucin-Degrading Bacterium. Int J Syst Evol Microbiol 2004, 54, 1476–, doi:10.1099/ijs.0.02873-0.

28. Laursen, M.F.; Larsson, M.W.; Lind, M.V.; Larnkjær, A.; Mølgaard, C.; Michaelsen, K.F.; Bahl, M.I.; Licht, T.R. Intestinal Enterococcus Abundance Correlates Inversely with Excessive Weight Gain and Increased Plasma Leptin in Breastfed Infants. FEMS Microbiology Ecology 2020, 96, fiaa066, doi:10.1093/femsec/fiaa066.

29. Jin, J.; Cheng, R.; Ren, Y.; Shen, X.; Wang, J.; Xue, Y.; Zhang, H.; Jia, X.; Li, T.; He, F.; et al. Distinctive Gut Microbiota in Patients with Overweight and Obesity with Dyslipidemia and Its Responses to Long-Term Orlistat and Ezetimibe Intervention: A Randomized Controlled Open-Label Trial. Frontiers in Pharmacology 2021, 12.

30. Ignacio, A.; Fernandes, M.R.; Rodrigues, V.A.A.; Groppo, F.C.; Cardoso, A.L.; Avila-Campos, M.J.; Nakano, V. Correlation between Body Mass Index and Faecal Microbiota from Children. Clinical Microbiology and Infection 2016, 22, 258.e1-258.e8, doi:10.1016/j.cmi.2015.10.031.

31. Zhuang, X.; Xiong, L.; Li, L.; Li, M.; Chen, M. Alterations of Gut Microbiota in Patients with Irritable Bowel Syndrome: A Systematic Review and Meta-Analysis. Journal of Gastroenterology and Hepatology 2017, 32, 38–, doi:10.1111/jgh.13471.

32. Jean, S.; Wallace, M.J.; Dantas, G.; Burnham, C.-A.D. Time for Some Group Therapy: Update on Identification, Antimicrobial Resistance, Taxonomy, and Clinical Significance of the Bacteroides Fragilis Group. Journal of Clinical Microbiology 2022, 60, e02361–20, doi:10.1128/jcm.02361-20.

33. Becker, H.E.F.; Jamin, C.; Bervoets, L.; Boleij, A.; Xu, P.; Pierik, M.J.; Stassen, F.R.M.; Savelkoul, P.H.M.; Penders, J.; Jonkers, D.M.A.E. Higher Prevalence of Bacteroides Fragilis in Crohn’s Disease Exacerbations and Strain-Dependent Increase of Epithelial Resistance. Front Microbiol 2021, 12, 598232, doi:10.3389/fmicb.2021.598232.

34. Chan, J.L.; Wu, S.; Geis, A.L.; Chan, G.V.; Gomes, T.A.M.; Beck, S.E.; Wu, X.; Fan, H.; Tam, A.J.; Chung, L.; et al. Non-Toxigenic Bacteroides Fragilis (NTBF) Administration Reduces Bacteria-Driven Chronic Colitis and Tumor Development Independent of Polysaccharide A. Mucosal Immunology 2019, 12, 177–, doi:10.1038/s41385-018-0085-5.

35. Wu, L.; Park, S.-H.; Kim, H. Direct and Indirect Evidence of Effects of Bacteroides Spp. on Obesity and Inflammation. Int J Mol Sci 2023, 25, 438, doi:10.3390/ijms25010438.

36. Blandford, L.E.; Johnston, E.L.; Sanderson, J.D.; Wade, W.G.; Lax, A.J. Promoter Orientation of the Immunomodulatory Bacteroides Fragilis Capsular Polysaccharide A (PSA) Is off in Individuals with Inflammatory Bowel Disease (IBD). Gut Microbes 2019, 10, 577–, doi:10.1080/19490976.2018.1560755.

37. Aggeletopoulou, I.; Kalafateli, M.; Tsounis, E.P.; Triantos, C. Exploring the Role of IL-1β in Inflammatory Bowel Disease Pathogenesis. Front Med (Lausanne) 2024, 11, 1307394, doi:10.3389/fmed.2024.1307394.

